# White-matter-microstructure-informed whole-brain models reveal localized excitation-inhibition imbalance in schizophrenia

**DOI:** 10.64898/2026.04.02.716059

**Authors:** Kailin Zhu, Georg Reich, Xiaojun Zhou, Trang-Anh E. Nghiem

## Abstract

Providing early diagnosis and personalized treatment for psychiatric disorders like schizophrenia remains challenging, due to important interpersonal differences and still elusive neuronal mechanisms. Whole-brain network models show promising results with clinical relevance for individualized treatment recommendations in neurological disorders. However, their applicability to psychiatry is still limited as models fail to account for inter-individual differences in the correlation structure of brain dynamics. What physiological mechanisms should models incorporate to better account for individual profiles of brain dynamics in schizophrenia patients and healthy controls? Our study compares various metrics of white matter structure and microstructure to inform connection weights between regions. To do so, we inferred regional parameters of whole-brain mean-field models with The Virtual Brain simulator to account for empirical functional connectivity from resting-state functional magnetic resonance imaging of schizophrenia patients and healthy controls. We found that using global fractional anisotropy or apparent diffusion coefficient of white matter fibers to inform the weights in neural mass models can drastically improve model performance. The data-model correlations of simulated and empirical data were significantly improved (from 0.2 to 0.7) over using the number or density of fibers as in many state-of-the-art methods. This approach allows us to uncover personalized maps of excitation-inhibition imbalance, hypothesized to underlie symptoms in schizophrenia. These maps prove meaningful in that they can predict diagnosis better than model-independent neuroimaging benchmarks. Our findings highlight the importance of white matter microstructure in whole-brain modeling. The novel white-matter-informed models reveal mechanisms that can cause altered brain dynamics in schizophrenia and could inform treatment in personalized psychiatry.

## 1 Introduction

Schizophrenia affects more than 23 million people worldwide [16]. Symptoms are wide-ranging across domains including hallucinations, delusions, cognitive control, and social interaction. Important inter-subject variability in symptoms and biomarkers has crucially hampered our ability to detect biomarkers of schizophrenia for early diagnosis as well as to develop widely effective treatments. Moreover, specifically targeting treatments to limit secondary effects is still challenging due to the fundamental gap in our knowledge of the cellular mechanisms causing altered brain function behind symptoms. What cellular mechanisms, in which brain regions, underlie altered brain dynamics in each schizophrenia patient? At the neuronal level, imbalance between excitation (E) and inhibition (I) due to loss of inhibitory conductance has been hypothesized to underlie symptoms [1]. As an alternative hypothesis, dysfunction of the dopamine system has been proposed - though recent evidence suggests a causal association between E/I imbalance and dopaminergic dysregulation in schizophrenia [6]. White matter structure and microstructure, as measured by diffusion-weighted magnetic resonance imaging (dwMRI), has also been observed to be altered schizophrenia [11]. However, it remains unclear whether cellular-level E/I imbalance or circuit-level white-matter could explain brain dynamical differences in schizophrenia. Further, it is still unclear which brain regions may be affected by these mechanistic alterations. The multi-domain nature of schizophrenia symptoms suggests that multiple brain regions may be affected, with distinct altered areas associated with different symptoms across patients. For instance, hallucinations may be caused by imbalance in sensory areas, with the most common hallucinations, auditory verbal i.e. hearing voices, possibly associated with E/I imbalance among language processing areas in the temporal lobe [8] and the salience network [10]. The default-mode network, associated with self-referential and social cognition, has also been found to be affected in schizophrenia among other disorders, including in its interactions with the salience network [9]. However, it remains difficult to map E/I balance at the individual level *in vivo*, as modern methods in magnetic resonance spectroscopy to probe E and I neurotransmitter concentrations need to be targeted to a single hypothesized region at a time.

To address this crucial challenge, personalized whole-brain models have emerged as a powerful paradigm to probe brain mechanisms behind brain dynamics at the individual level [7, 20, 15, 4]. Briefly, whole-brain models simulate brain activity based on differential equations for each region, with connection weights and delays between regions informed by individualized white-matter data. Whole-brain models have proven clinically relevant for E/I imbalance disorders like epilepsy, by helping localize the epileptic focus in each patient to guide surgery [7]. However, inferring parameters like inhibitory conductance, controlling E/I imbalance, at the regional level has remained computationally challenging, until new developments introduced E-to-E and E-to-I inter-regional coupling inference [15]. Yet, these novel tools remain to be tested in their ability to account for individual brain dynamics in patient populations. Moreover, decades of literature on whole-brain modeling have never systematically investigated the effects of white-matter-microstructure in shaping FC. In most models, when specified, the number or density of fibers is used to inform connection weights between regions. However, microstructural white-matter properties like generalized fractional anisotropy (gFA) and apparent diffusivity coefficient (ADC) may also modulate inter-regional interactions, which whole-brain modeling has never accounted for.

In this manuscript, for the first time, we systematically compare metrics of white matter structure and microstructure to inform personalized simulations of brain activity in schizophrenia and controls. To do so, we infer regional parameters of whole-brain mean-field models with The Virtual Brain (TVB) to account for individual functional connectivity (FC) from resting-state functional magnetic resonance imaging (fMRI) data. First, we compare models informed with different white-matter metrics in their fit to empirical FC. Then, we uncover the relevant aspects of white-matter data to capture FC, by shuffling white-matter matrices across regions and participants and testing whether data-model correlations worsen. Next, we reveal which regions are affected by E/I imbalance in schizophrenia by investigating inferred E/I parameter maps. Finally, we demonstrate that inferred E/I maps are meaningful by leveraging them for diagnostic classification and comparing accuracy with metrics independent of our model. Our approach provides a white-matter-microstructure-informed platform to model brain activity at the individual level in health and pathology, allowing us to introduce and validate tools to map E/I balance and derive mechanistic inductive biases supporting machine-learning-based diagnostics.

## 2 Materials and Methods

### Data

The dataset used for this study [19] consists of structural and functional connectomes from 27 (age 41 ± 9.6) schizophrenic patients and 27 matched healthy controls (age 35 ± 6.8 years). All patients met the DSM-IV criteria for a diagnosis of schizophrenic or schizoaffective disorders and were under medication. All data is from eyes-open resting-state fMRI. A Desikan-Killiany parcellation into 83 cortical and subcortical areas was applied to the data. In one replication analysis, we also consider a parcellation into 129 areas. For each parcellation, considered white matter metrics quantifying structural connectivity (SC) included the number of fibers i.e. streamlines between regions, the density of streamlines normalized by their length and regional surface area, as well as the gFA and average ADC defined by the anisotropy of inter-regional streamline directions and water diffusivity respectively. Functional connectivity was computed as pairwise Pearson correlation between the fMRI signal of regions, after detrending, smoothing, removal of artifacts, and exclusion of the first four time points.

### TVB simulations

We used The Virtual Brain (TVB) simulator to construct personalized whole-brain network models and simulate regional synaptic activity and functional connectivity (FC) [4]. Each brain region *i* is modeled as an E-I population governed by a reduced Wong–Wang neural mass model [2, 15], capturing slow NMDA-mediated synaptic dynamics relevant for resting-state fMRI and widely used in whole-brain functional connectivity modeling. The E and I firing rates 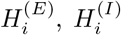 and corresponding synaptic gating variables 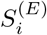 and 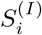 are given by

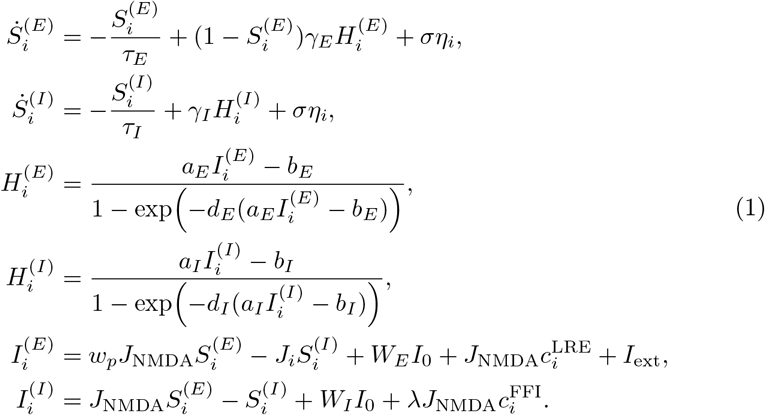

Inter-regional couplings are defined by

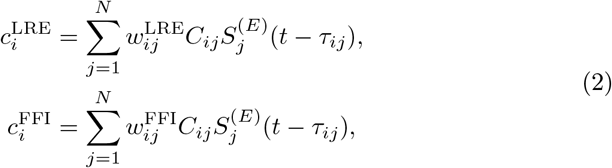

The parameters *τ*_(*E,I*)_ and *γ*_(*E,I*)_ control synaptic time scales and saturation, while *J*_NMDA_ and *w*_*p*_ determine excitatory coupling and local recurrence, and *J*_*i*_ denotes local inhibitory feedback. The inputs 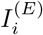 and 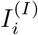 combine local excitation–inhibition, background drive *W*_(*E,I*)_*I*_0_, external input *I*_ext_, and delayed inter-regional coupling. Between regions, coupling strength and delays are given by the normalized empirical structural connectivity is *C*_*ij*_ and delays *τ*_*ij*_ derived assuming a constant conduction speed of 3.0 m*/*s, while 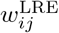 and 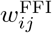 denote long-range excitation and feedforward inhibition weights from region *j* to *i*. The term *ση*_*i*_ represents additive Gaussian noise.

### TVB inference

To maximize the Pearson correlation between simulated and empirical FC, we infer regional and inter-regional parameters *J*_*i*_, *w*^LRE^, and *w*^FFI^ at the individual level. Regional parameters are adjusted using feedback inhibition control [2]. At each optimization step, regional I feedback parameters *J*_*i*_ are updated according to 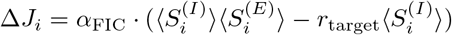, with ⟨*S*^(*E*)^⟩ the synaptic gating activity variable for the excitatory population. This ensures that the average firing rate over time does not diverge or saturate in each region. Simultaneously, the inter-region coupling weights to E and I populations, *w*^LRE^ and *w*^FFI^, are adjusted by update rules based on the difference between simulated and empirical FC following 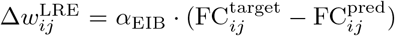 · RMSE_*i*_ and 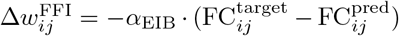 [12]. Simulated FC was computed by hemodynamic response function convolution [12].

### White-matter matrix shuffling

To interpret our findings, we use a permutation approach by randomly shuffling first the labels of the participants and then, the weights of the white-matter SC matrices. For the first test, we randomly permute participant’s labels such that each person’s FC is simulated using another person’s SC matrix. In the second part, we shuffle across the regions in each individual to test whether regional specificity has influence on model fit of empiral FC. To do so, we randomly shuffle region labels, using the same shuffled list of labels across participants. Next, the SC matrix for each participant was rearranged according to the shuffled region labels and used to fit their FC matrix. This ensures that overall graph properties are preserved but the regional specificity was destroyed. For both shuffling analyses, models are inferred again with these different SC matrices.

### Diagnostic classification

To study whether inferred model parameters were meaningful and useful for diagnosis, we use stochastic gradient descent (SGD) classifiers from the python *scikit-learn* library. As inputs to the classifiers, we provide vectors of model-inferred *J*_*i*_, 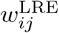, and 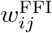parameters for each participant. Using these inputs, the classifiers are trained to discriminate between schizophrenia and control diagnostic labels. The dataset is randomly split into training and test sets of equal size. Performance was then evaluated using test accuracy across *n* = 100 random splits. For comparison, model-independent inputs consisting of concatenated FC and SC matrices are given to the classifiers. We use the hinge, log loss, modified Huber, and perceptron loss functions. The maximum iteration number is set to 5000 and the stopping criterion is given by *tol* = 5 * 10^−3^. To prevent overfitting, L2 regularization is applied.

### Statistical testing

All p-values are obtained from non-parametric Wilcoxon sign-ranked tests. We use the Holm-Sidak correction for multiple comparisons over metrics, pair of metrics, or loss functions wherever applicable.

## 3 Results

### 3.1 Informing whole-brain models with white matter ADC and gFA drastically improves data-model fit

To systematically test to what extent integrating white-matter microstructure into whole-brain modeling helps account for the data, we simulate brain activity using, as inter-regional weight matrices *C*_*ij*_, successively only the number of fibers, only the density of fibers, only the ADC, and only the gFA. All other parameters are either kept identical across simulations or inferred at the individual level for direct comparison purposes. This procedure is repeated for the data of each schizophrenia patient and healthy control: for each individual, we carried out simulations successively with each of their white-matter metrics as weight matrices, and inferred model parameters to match their personal FC matrix. We find that data-model correlations are considerably larger (*r* ≈ 0.7) for models informed by ADC and gFA compared to the state-of-the-art models informed by the number and density of fibers (*r* ≈ 0.2) (Table 1). This difference is statistically significant across both schizophrenia and control participants (\*\*\**p* <10^−8^ for all metric pairs except between ADC and gFA). No significant differences are found between data-model correlations in patients vs controls (*p* > 0.05). Results are replicated across both 83-region and 129-region brain parcellations. The results emphasize the role of white-matter microstructure in sculpting restingstate FC and the importance of informing whole-brain models with ADC or gFA metrics to capture brain activity in both health and schizophrenia.

**Table 1:**
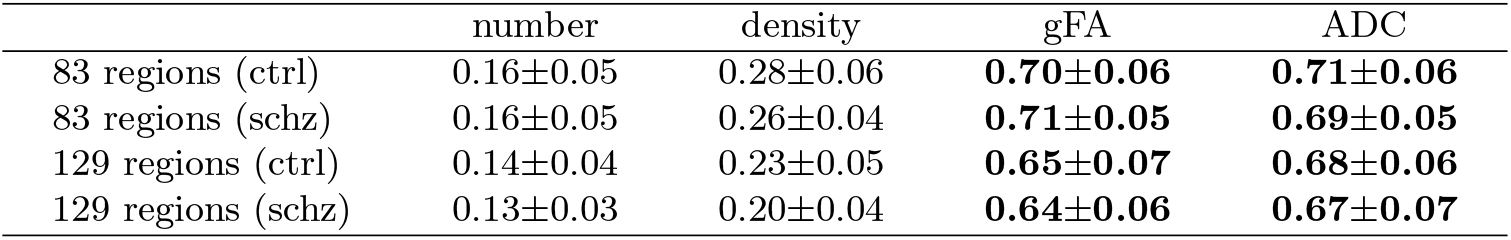
E/I tuning model Pearson FC correlation (mean ± std)

|  | number | density | gFA | ADC |
| --- | --- | --- | --- | --- |
| 83 regions (ctrl) | 0.16±0.05 | 0.28±0.06 | **0.70±0.06** | **0.71±0.06** |
| 83 regions (schz) | 0.16±0.05 | 0.26±0.04 | **0.71±0.05** | **0.69±0.05** |
| 129 regions (ctrl) | 0.14±0.04 | 0.23±0.05 | **0.65±0.07** | **0.68±0.06** |
| 129 regions (schz) | 0.13±0.03 | 0.20±0.04 | **0.64±0.06** | **0.67±0.07** |

### 3.2 The regional specificity, but individual specificity of ADC and gFA graphs, are relevant for data-model fit

To uncover the aspects of ADC and gFA supporting model fit, we perform a permutation analysis across all participants as well as brain regions. Surprisingly, for ADC-informed models, shuffling the white-matter matrices across the 54 participants yields no significant difference for any metric in model performance (Fig. 2, *p* = 0.7). The same non-significant effects are observed upon shuffling gFA, number, and density matrices (*p* > 0.05 for all metrics). In contrast, shuffling across brain regions in each individual and thus, disrupting regional specificity in structural connectivity matrices, we observe a significant reduction in model performance. This is the most prominent in ADC, number of fibers, and density (\*\*\**p* <0.0001) for both control and schizophrenia groups with both metrics. However, for the models informed by gFA, inter-regional shuffling worsens fit significantly for schizophrenia patients but not controls (\**p* = 0.02 and *p* = 0.2 respectively). The results show that regional specificity of ADC and gFA matrices matters but not individual specificity.

**Fig. 1:**
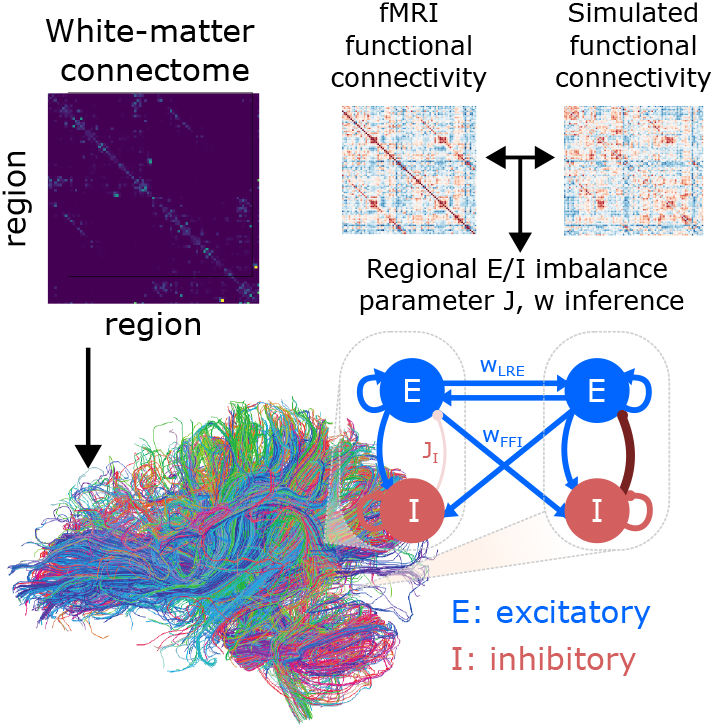
Diagram of the model and inference procedure. After obtaining the individual connectome from white-matter data, we construct a personalized whole-brain dynamical model using reduced Wong–Wang excitatory– inhibitory equations for each brain region. Then by fitting simulated FC to empirical FC, we infer regional and inter-regional E/I parameters for every individual.

**Fig. 2:**
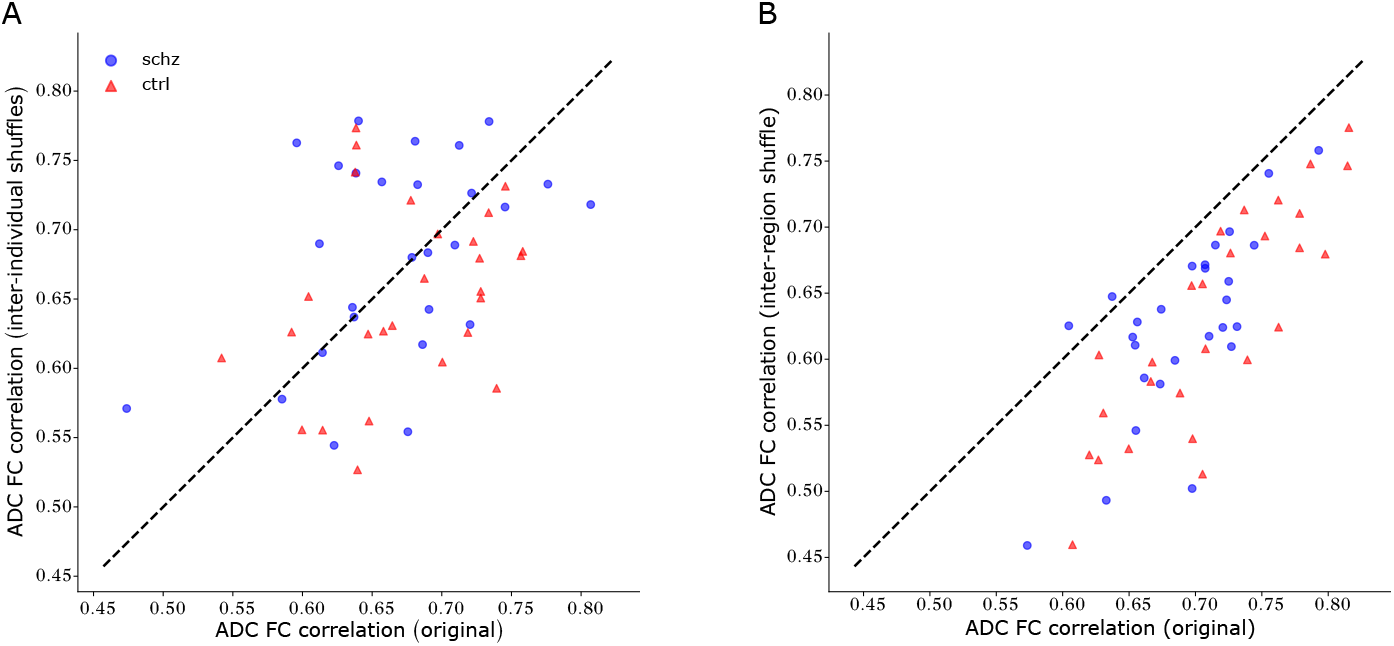
Correlations between empirical and simulated FC, informed by ADC, for original vs **(A)** inter-individually and **(B)** inter-regionally shuffled weights. Each point represents one subject. The dashed diagonal line is the identity. Points below the diagonal indicate worsened model performance after shuffling.

### 3.3 Inferred E/I parameter maps reveal putative imbalance among posterior cingulate and paracentral regions

To localize regions of E/I imbalance, we further investigate the inferred *J*_*i*_ parameters and their difference between controls and schizophrenia, averaged across participants in each group. Regions that appear among the ten with the largest difference for both ADC and gFA included the left and right posterior cingulate cortex (PCC), an area of the default-mode network, and paracentral region, a sensorimotor integration area overlapping with the language processing system, which seems robust across all metrics except the number of fibers for the posterior cingulate cortex (Fig. 3). Notably, the largest difference observed for gFA was in the accumbens, followed by the isthmus cingulate, which are not found for other metrics. For ADC, the most prominent difference is observed in the PCC. The results show that the PCC and paracentral regions are the most consistently affected by E/I imbalance in schizophrenia according to our model.

**Fig. 3:**
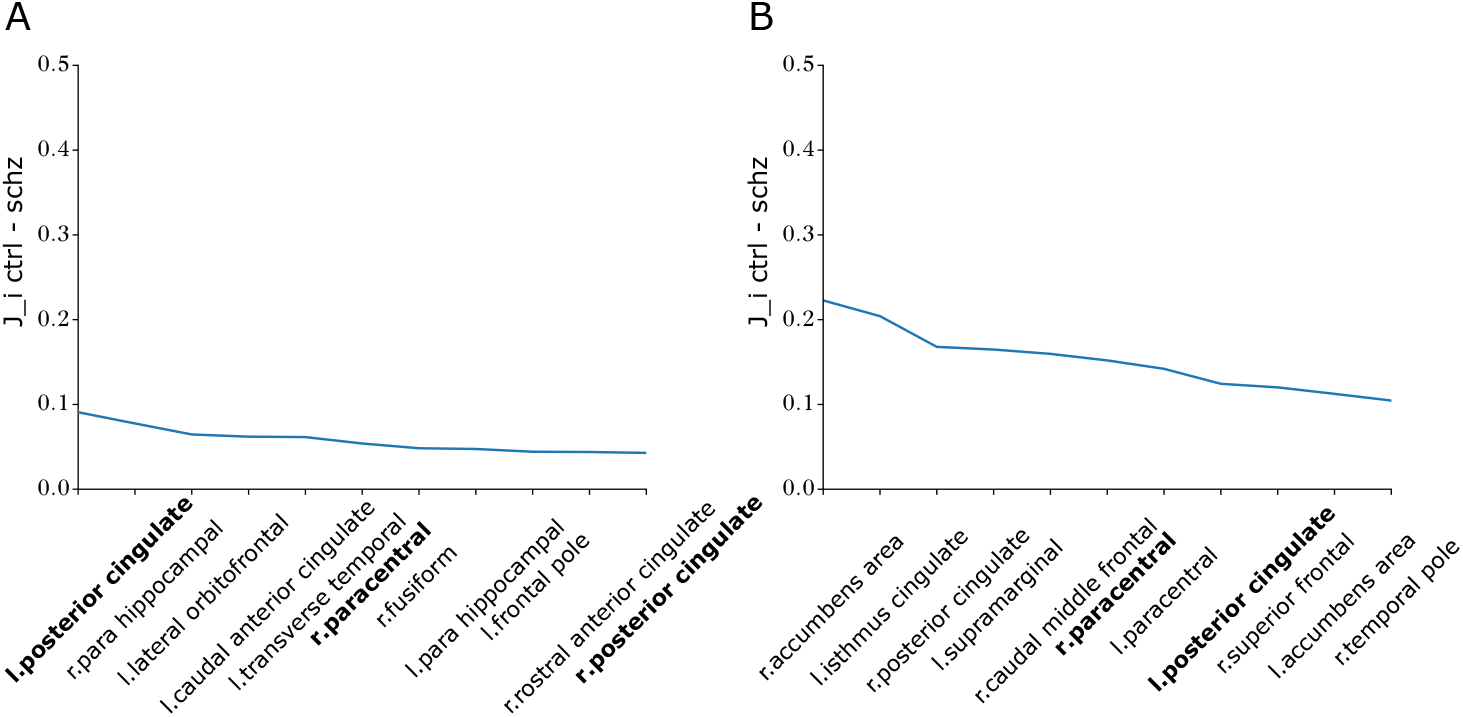
Regional *J*_*i*_ differences between control and schizophrenia groups. The top ten regions are ranked in decreasing order of *J*_*i*_ differences for (**A**) ADC - and (**B**) gFA - informed models. Regions appearing in both are highlighted in bold.

### 3.4 Using inferred E/I maps tends to improve diagnostic classification over using model-independent metrics

To investigate whether E/I maps can underpin symptoms and test their utility for diagnosis, we compare diagnostic classification from model-inferred E/I balance parameters vs empirical FC and SC matrices. Using the wFFI to inform the classifier yields a mean accuracy of 0.7 for ADC across random data splits. This significantly exceeds the test scores obtained using combined empirical FC and SC data which accounts for a test score of 0.6 (\*\*\**p* <0.001). The findings are robust across loss functions (Fig. 4). The results support that E/I map inference amplifies clinically relevant dimensions of SC and FC data, such that E/I maps can enable diagnostic classification at benchmark-comparable levels.

**Fig. 4:**
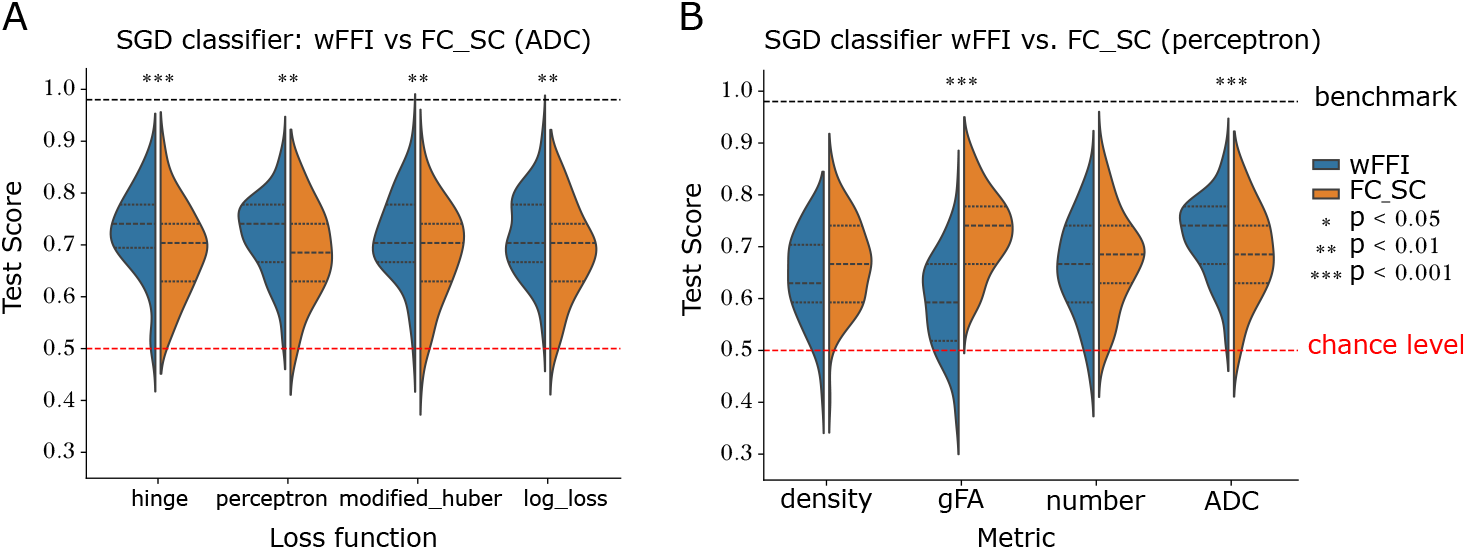
Diagnostic classification accuracy using E/I balance model parameters. (**A**) Accuracy across different loss functions, comparing SGD classifier informed by simulated wFFI and empirical FC and SC. The models were informed by ADC. The red line indicates chance level and the black line the benchmark performance in the field [14]. Each plot shows the distribution of test scores for 100 splits. **(B)** Accuracy across different metrics for perceptron loss.

## 4 Discussion

In this work, we compared whole-brain models informed by various white-matter metrics in their ability to account for resting-state fMRI in schizophrenia and controls. We found that models informed by gFA and ADC microstructural metrics consistently outperformed models informed by state-of-the-art number and density of fibers at capturing empirical FC. The regional specificity, but not the individual variability of white-matter matrices was crucial to reproduce subject FC in simulations, which suggests that in our models, E/I imbalance accounts for most of inter-subject and schizophrenia-control differences. E/I imbalance maps revealed reduced inhibition in the PCC and paracentral areas of the default-mode and sensorimotor networks in schizophrenia. Maps were also meaningful in that they were relevant for diagnostic classification. The findings support that whole-brain models informed with white-matter microstructure can capture the structure of fMRI correlations and meaningfully map E/I imbalance underlying brain dynamical differences and potentially symptoms in schizophrenia.

First, we find that informing models with gFA or ADC yields significantly improved data-model correlations (*r* ≈ 0.7) compared to using the number or density of fibers as commonly used in the literature (*r* ≈ 0.2). Here, our findings contribute a systematic comparison of these metrics to the literature, when different past works may have used different white-matter metrics to fit various datasets. A correlation between empirical and simulated FC of *r* ≈ 0.7 improves on even recent works [13]. In particular, fitting regional and inter-regional inhibition parameters supports model performance, consistent with published work that found up to *r* ≈ 0.7 by inferring regional inhibition [20], including on the same data or with the same pipeline [3, 12]. Together with our results, this supports the importance of regional E/I balance in shaping FC. To our knowledge, our research is the first to apply this approach to patient data, allowing us to observe that the methods are also relevant to mapping pathological E/I imbalance towards uncovering root causes of symptoms and guiding treatment.

Second, we reveal that shuffling white-matter matrices across participants did not significantly worsen model performance. Even shuffling between participants with schizophrenia and controls did not worsen fit to data, surprisingly as white-matter microstructure was shown to be affected in schizophrenia [11]. Our results suggest, by contrast, that resting-state dynamical alterations in schizophrenia are not explained by underlying white-matter microstructure. However, shuffling between regions gave rise to small, although significant, changes in data-model fit. This is due to inter-regional E/I balance parameters playing a comparatively crucial part in shaping FC, with the potential confound of overfitting due to heavy parametrization of inter-regional E-to-E and E-to-I weights. Taken together, these results open doors to using white-matter microstructure from one dataset to fit FC from other data where white matter data are unavailable, hence increasing the applicability of personalized whole-brain modeling beyond the state of the art where individual diffusion data is needed.

Our models are heavily parametrized in that inferred regional and inter-regional maps importantly contribute to model-data fit over white-matter specifics. Hence, it is crucial to ensure that inferred E/I imbalance parameter maps are meaningful. A first way to tackle this validation question is to investigate regions uncovered by the model as affected by E/I imbalance in schizophrenia. This approach revealed that the PCC and paracentral areas showed on average reduced inhibition in schizophrenia. Dynamical differences consistent with E/I imbalance were also observed in the PCC in autism spectrum disorder, which shares social interaction symptoms with schizophrenia [17]. The PCC is also associated with autobiographical memory, potentially related to delusion symptoms domains. The paracentral area is a sensorimotor region among areas involved in language and salience processing, relevant to auditory verbal hallucinations and found to be affected in schizophrenia [18]. Thus, the regions found by our model are in line with the literature, which supports the validity of our method. Our model produces a putative explanation for dynamical differences in these regions, by suggesting they could be due to reduced inhibition compared to excitation.

Another way to validate our approach is to use model-derived E/I balance parameters to predict diagnosis, as the model was not informed with diagnostic data. We find that the inferred regional and inter-regional parameters controlling E/I balance can outperform model-independent SC and FC data for diagnostic classification. Here, the ADC appears from our analyses as a white-matter metric able to both capture resting-state FC and diagnostic-relevant information. State-of-the-art diagnostic prediction methods perform similarly or better than E/I balance maps [18, 5, 14], which could be because they take into account dynamical aspects of fMRI signals beyond average FC over entire time series. Thus, future work inferring E/I balance maps as dynamical, time-varying parameters from patient data may yield diagnostic improvements. Overall, our method may be understood as a way to analyze data, combining information from SC, FC, and E/I balance biophysics to generate inductive biases for clinical machine learning tasks like diagnostics. By amplifying data directions relevant that are diagnostically relevant and associated with hypothesized causal mechanisms, our white-matter-microstructure informed modeling may be useful to replace or add to existing inductive biases for clinical machine learning tasks.

Importantly, our work has several limitations. First, we provide a proof of prin-ciple on a small dataset of 27 schizophrenia patients and 27 control participants [19]. It remains to be seen if results generalize to larger and more heterogeneous cohorts such as including early psychosis. Investigation in larger cohorts would also enable the clustering of patients into disorder subtypes potentially corresponding to different E/I imbalance maps. Moreover, the effects we identify are observed in resting state and do not necessarily generalize to brain function in tasks tied to symptoms. While we chose to focus on resting-state data as it is currently the most readily available in public datasets from schizophrenia patients, extensions of this approach to tasks tied to symptoms may provide more insights into the causal role of E/I balance in altered brain function.

In conclusion, we developed personalized and white-matter-microstructure informed models of brain dynamics in health and schizophrenia. Our results provide the first systematic investigation of the role of white matter in whole-brain modeling, emphasizing the importance of ADC and gFA metrics. This approach reveals, for the first time, maps of E/I imbalance in a patient population, with reduced inhibition in distinct brain regions across functional networks. Inferred imbalance maps are meaningful in that they can support clinical machine learning tasks like diagnostic classification. The findings create a fundamentally novel bridge between cellular-scale E/I imbalance mechanisms hypothesized in schizophrenia and brain-wide dynamics that can be recorded *in vivo* in patients. Personalized white-matter microstructure informed models could therefore be relevant to simulate disorder progression for early diagnosis and to optimize intervention protocols toward individualized treatment recommendations.

